# An Open–Access Simultaneous Electrocardiogram and Phonocardiogram Database

**DOI:** 10.1101/2021.05.17.444563

**Authors:** Arsalan Kazemnejad, Peiman Gordany, Reza Sameni

**Affiliations:** School of Electrical & Computer Engineering, Shiraz University, Shiraz, Iran; Department of Biomedical Informatics, Emory University School of Medicine, GA 30322, US

**Keywords:** Electrocardiogram, Phonocardiogram, Stress-test

## Abstract

The electro-phono-cardiogram (EPHNOGRAM) project focused on the development of low-cost and low-power devices for recording simultaneous electrocardiogram (ECG) and phonocardiogram (PCG) data, with auxiliary channels for capturing environmental audio noise, which could be used for PCG quality enhancement through signal processing. The current database, recorded by version 2.1 of the developed hardware, has been acquired from 24 healthy adults aged between 23 and 29 (average: 25.4 ± 1.9 years) in 30 min stress-test sessions during resting, walking, running and biking conditions, using indoor fitness center equipment. The dataset also contains several 30 s sample records acquired during rest conditions. This data is useful for simultaneous multi-modal analysis of ECG and PCG. It provides interesting insights into the inter-relationship between the mechanical and electrical mechanisms of the heart, under rest and physical activity. The database is provided online on PhysioNet.

## I. Introduction

Cardiac auscultation is one of the oldest and most basic methods of cardiac function assessment. Even in the modern cardiac monitoring and imaging era, the technique remains popular among clinicians, as a preliminary step for screening basic cardiac anomalies. Despite its long history, the visualization, analysis and interpretation of the audio signals acquired from the heart by the phonocardiogram (PCG) is not so common in clinical training. Therefore, PCG-based diagnosis is less common than its electrical counterpart, the electrocardiogram (ECG). Nevertheless, with recent developments in mobile-health and tele-monitoring, the PCG and ECG are again under the spotlight, as low-cost complementary modalities for monitoring the mechanical and electrical functions of the heart [1]. While most research in this domain have used separate sessions of ECG and PCG acquisition (which are useful for consistent cardiac anomalies), they do not provide a beat-wise insight into the two cardiac modalities and the interrelationships between the electro-mechanical functions of the heart.

The electro-phono-cardiogram (EPHNOGRAM) project conducted at the Signal Processing Center of Shiraz University (Shiraz, Iran) focused on the development of low-cost and low-power devices for recording simultaneous ECG and PCG [2]. As part of this research a hand-held system was designed and used to acquire a dataset consisting of simultaneous ECG and PCG of healthy adults in a stress-test study. The developed hardware has various features including auxiliary channels for capturing environmental audio noise, which could be used for PCG quality enhancement through signal processing.

The gathered database is provided online on PhysioNet [3], and the source codes for reading and analyzing the data are available in the *Open-Source Electrophysiological Toolbox* (OSET) [4].

In Section II, the architecture of the designed system is detailed. The stress-test dataset and its acquisition protocol is detailed in Section III. As proof of concept for the potential outcomes of this study, some of the basic analysis on these datasets are presented in Section IV. We have developed a robust and accurate algorithm for calculating the heart-rate (HR) from the ECG and PCG. The motivation is to show that while the overall trend of the HR time-series obtained from both modalities are identical, there are minor differences between them, which reflect the differences between the mechanical and electrical systems of the heart, during physical activities.The paper is concluded with a discussion and future perspectives for simultaneous ECG-PCG analysis.

## II. The EPCG device

The EPCG device designed for this study includes circuitry for three-lead ECG, two digital stethoscope channels for PCG acquisition and two auxiliary channels to capture the ambient noise, as shown in Fig. 1. The auxiliary channels are used for digital active noise cancellation (ANC) from the primary PCG channels. The analog signals are filtered by an anti-aliasing analog filter and sampled at 8 kHz with a resolution of 12-bits (with 10.5 effective number of bits) and transferred to an on-board low-power microcontroller for minimal preprocessing and registration on a Secure Digital (SD) memory.

**Fig. 1.**
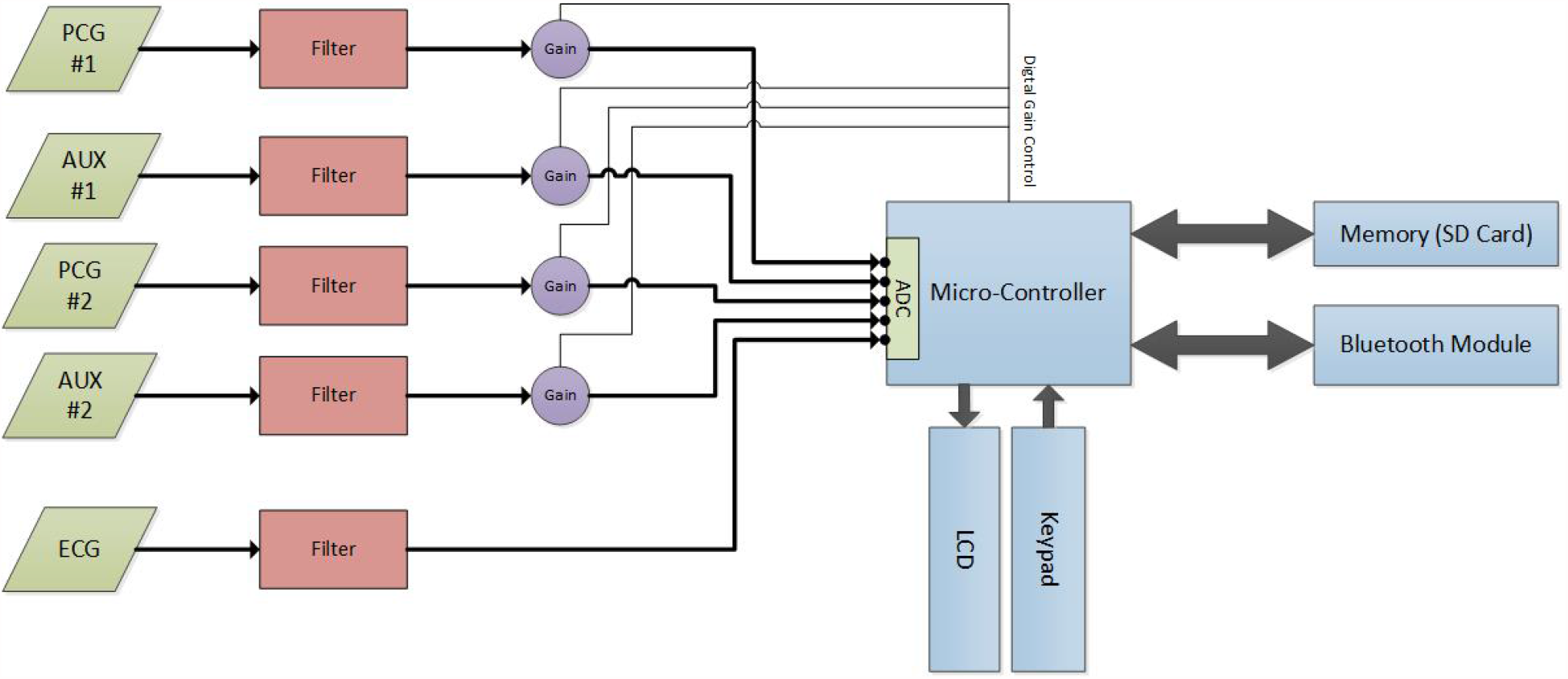
The ECG-PCG acquisition system block diagram

The device has an LCD and a keypad for basic file naming and controlling the recording duration. Since the quality of the ECG highly depends on the connection of the chest leads, the device provides an online PC-based signal preview feature via Bluetooth, to prevent low-quality signal recordings due to loose body contact.

The front-end anti-aliasing and baseline wander rejection filter consists of a first-order passive high-pass filter with a –3 dB cutoff frequency of 0.1 Hz, followed by an active 5th order low-pass Butterworth filter, which form bandpass filters that cover the major ECG and PCG bandwidths. For the ECG front-end, the upper –3 dB cutoff frequency was set to 150 Hz, with 30 dB of attenuation at 1 kHz and a 30 dB gain in the passband^1^. For the PCG channels, the same active filter topology was used, but with an upper cutoff frequency of 1 kHz, 30 dB of attenuation at 5 kHz, and a passband gain of 5 dB. As a result, the ECG and PCG channel filter attenuations are respectively 90 dB and 25 dB in the Nyquist frequency of the digital front-end ADCs (4 kHz), which are practically sufficient to avoid aliasing. Note that although the front-end filters are nonlinear phase, the group delays are rather constant over the passbands of the front-end filters. Additional filtering, including powerline cancellation (50 Hz for the current database) is performed in the digital domain.

During the prototyping process of the device, various configurations of the hardware and several hand-made stethoscopes were developed and tested. The hardware design Version 2.1 was the latest of several progressive improved redesigns, in which the auxiliary channels were added (useful for digital active noise cancellation algorithms) and the signal quality was significantly improved through the redesign of the analog front-end and the printed circuit board (PCB).

For the stethoscopes, the objective was convert low-price off-the-shelf stethoscopes into high-quality digital ones with minimal engineering effort and by using advances signal processing. The different designs that were prototyped and tested include: 1) embedding microphones inside standard stethoscope chest-piece (directly under the diaphragm); 2) embedding two microphones inside a standard microphone chest-piece (back-to-back, one facing the diaphragm, the other one facing the bell hole); 3) embedding microphones at the end of the tubing (at the junction of the ear tubes); 4) embedding microphones inside the tube, a few centimeters after the stem, plus auxiliary microphones inside the device case, for picking environmental noises. The latter configuration, as demonstrated in in Fig. 2, was found to be the most robust and resulted in high quality signals, with various advantages:

1. Arbitrary stethoscope diaphragms can be used with this device and modifications are only applied to the tubing section.
2. The acoustic frequency response of the mechanical parts of the chest-piece (diaphragm and bell) is not altered. In fact, the chest-piece is the most sophisticated piece of a stethoscope, in terms of its frequency response and the impact of its shape on the quality of the PCG. By embedding the microphone inside the tube (after the stem), the frequency response of the PCG undergoes the minimal change, compared to what clinicians hear with a classical stethoscope.
3. Along the stethoscope tube, the transfer function of the audio channel is approximately linear. Therefore, the audio quality is less susceptible to the precise position of the microphone (in orders of millimeters) inside the tube, which is an important factor for mass production.
4. The auxiliary channels uniquely pick the environmental sounds, which can be used in the software to implement adaptive noise cancellers or advanced source separation algorithms.

**Fig. 2.**
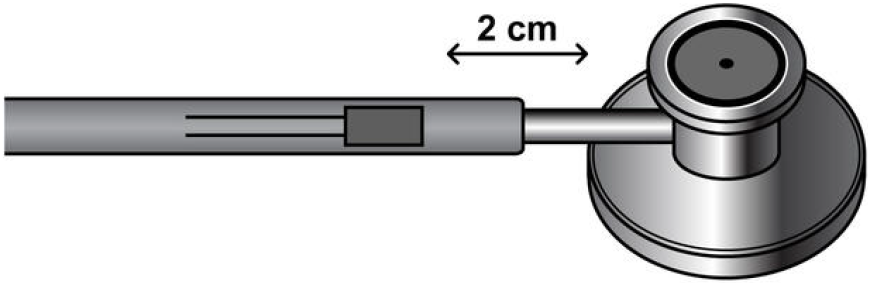
A schematic of the designed stethoscope with a microphone embedded in the tubing

It should be noted that various versions of this stethoscope were built, using capacitive, piezoelectric and MEMS microphones. The datasets introduced in the sequel was acquired by the capacitive microphone version of the stethoscope.

## III. The database

The acquisition of signals with the developed device was approved by the Biomedical Engineering Review Committee (IRB equivalent) of Shiraz University School of Electrical and Computer Engineering. A total number of 24 male subjects aged between 23 and 29 (average: 25.4 *±* 1.9 years) attended the study. The individuals gave informed consent to participate in the study.

The acquired dataset consists of 69 simultaneous ECG and PCG recordings, each with a duration of 30 seconds (8 records) and 30 minutes (61 records), acquired synchronously from a three lead ECG and a single PCG stethoscope. Each volunteer performed a specific task once (as detailed in the sequel). In a few cases where the data quality was poor (due to electrode/stethoscope detachment and analog frontend saturation), the test was repeated to obtain acceptable data. However, even the poor quality samples have been included in the dataset for noise research purposes and labeled as low quality in the spreadsheet accompanying the dataset. In addition to the main PCG channel, for some subjects the auxiliary audio channels PCG2, AUX1, and AUX2 were recorded for audio processing research purposes; although these auxiliary channels are mostly at quantization noise level for the majority of the recorded sessions.

Cardiac auscultation is commonly performed from four major chest areas. The Mitral valve (M) sound is heard better at the distal end of the heart, anatomically landmarked between the fifth and sixth ribs on the body surface. The Tricuspid valve (T) sound is heard well on the left side of the heart, between the fourth and sixth ribs. Therefore, we chose the location between the Tricuspid and Mitral landmarks to record the heart sound, to obtain both the first and second heart sounds to a good extent, as shown in Fig. 3. The placement of the three ECG leads are also shown in Fig. 3.

**Fig. 3.**
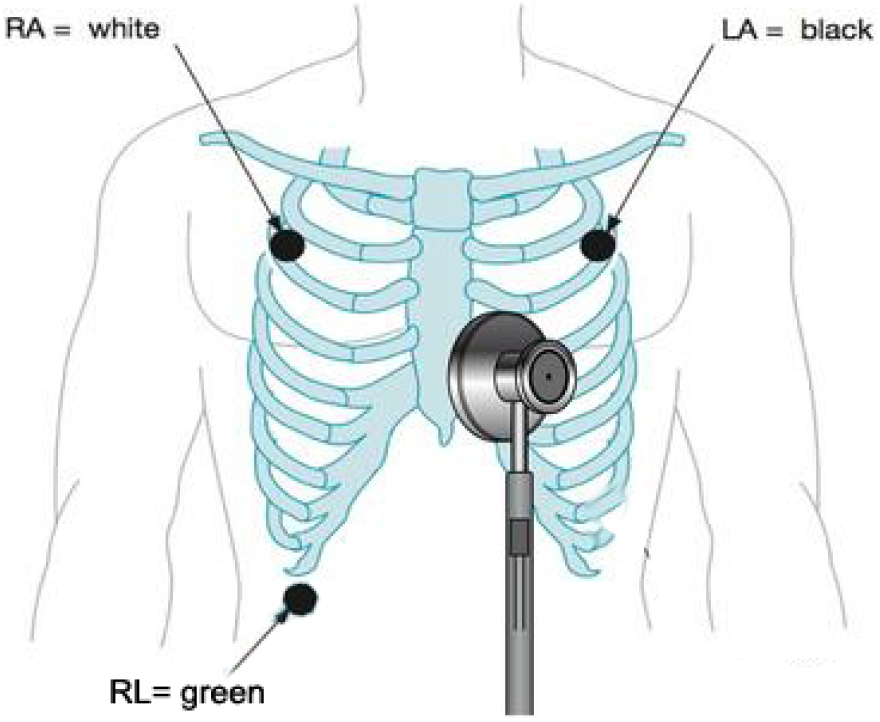
ECG lead configuration and PCG stethoscope position for Dataset1

### A. Acquisition protocol

The 30 s records were recorded during the development phase of the device and the participants were seated in an armchair during their acquisition.

The 30 minutes records (62 records) were acquired in an indoor sports center. A structured interview determined that the participants were in good physical condition and none reported symptoms of autonomic or cardiovascular disorder. Each subject participated in one or a number of the physical scenarios, detailed below. Accordingly, ten subjects participated in the first scenario. Five of the participants of the first scenario did not attend the rest of the stages. With the addition of six new volunteers, a total number of eleven subjects contributed in each of the three other scenarios. Only in the bicycle exercise stress-test (Scenario D), two volunteers failed the test due to physical fatigue. In order to prepare for each test, volunteers avoided eating food, drinking caffeine, alcoholic drinks and smoking for three hours before the test. But they were permitted to drink water regularly.

The four tested scenarios are as follows:

1. *Scenario A (rest condition):* The participant laid horizontally on a bed in a quiet room while ECG and PCG signals were recorded for thirty minutes.
2. *Scenario B (walking condition):* The participant walked on a treadmill at a constant speed of 3.7 km/h at an inclination angle of one degree. The process lasted thirty minutes at constant speed and slope.
3. *Scenario C (treadmill stress-test):* The modified Bruce protocol was used for the treadmill stress-test [6], as shown in Table I. This modified stress-test started at a lower workload as compared to the standard test. The test lasted thirty minutes and the increase in speed and treadmill slope continued until the subjects reached excessive fatigue, excessive heart-rate or chest pain. Whenever the subjects reached this (subject dependent) extreme point, in order to avoid a sudden decrease in the heart-rate, the speed was gradually decreased to 3 km/h and the slope was decreased to horizontal position. After a few minutes of walking in this state, the participant sat in a chair until the end of the test. The signals were acquired up to the end of the test.
4. *Scenario D (bicycle stress-test):* The bicycle stress-test protocol is detailed in Table II. The test lasted thirty minutes. The participant first rested for two minutes on a stationary exercise bicycle without pedaling, then started pedaling at a workload of 25 Watts/min. At the beginning of the test, according to Table II, the external load was gradually increased and the subject was asked to match the power consumption to the power specified in the standard test. Note that it is practically not possible to keep the pedal speed at a constant value. Therefore, the power consumption is only an approximation of the desired power consumption. The participants continued the test until they reached excessive fatigue, excessive heart-rate, or chest pain. Next, in order to prevent a sudden decrease in the heart-rate, the external load level was gradually decreased to 25 Watts/min. The subject remained in this state for several minutes, then rested on a stationary bicycle until the end of the test period, while the signals were continuously recorded.

**TABLE I.**
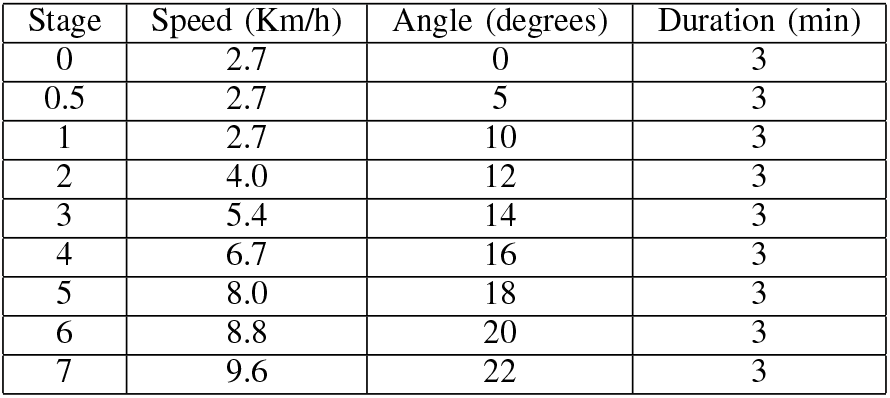
Modified Bruce treadmill stress-test protocol

| Stage | Speed (Km/h) | Angle (degrees) | Duration (min) |
| --- | --- | --- | --- |
| 0 | 2.7 | 0 | 3 |
| 0.5 | 2.7 | 5 | 3 |
| 1 | 2.7 | 10 | 3 |
| 2 | 4.0 | 12 | 3 |
| 3 | 5.4 | 14 | 3 |
| 4 | 6.7 | 16 | 3 |
| 5 | 8.0 | 18 | 3 |
| 6 | 8.8 | 20 | 3 |
| 7 | 9.6 | 22 | 3 |

**TABLE II.** Bicycle exercise stress-test protocol

| Stage | Power consumption (Watts/min) | Duration (min) |
| --- | --- | --- |
| 0 | 0 | 2 |
| 1 | 25 | 2 |
| 1 | 25 | 2 |
| 2 | 50 | 2 |
| 3 | 75 | 2 |
| 4 | 100 | 2 |
| 5 | 125 | 2 |
| 6 | 150 | 3 |
| 7 | 175 | 2 |
| 8 | 200 | 2 |
| 9 | 225 | 2 |
| 10 | 250 | 2 |
| 11 | 275 | 2 |
| 12 | 300 | 2 |

### B. Data files on PhysioNet

The complete dataset is available online on PhysioNet [3]. The data files are presented in both MATLAB (`ECGPCG00XY.mat` where `XY` = 01,…, 69) and WFDB (`ECGPCG00XY.dat` and `ECGPCG00XY.hea`) formats, with identical base names. The MATLAB files are in double-precision floating point format. Each file was covered into 16-bit WFDB format by using the `mat2wfdb.m` function from the WFDB Toolbox [7], [8]. The accuracy of conversion between MATLAB and WFDB formats was assessed per file and per channel, by comparing the signal-to-noise ratio (SNR) of the original MATLAB files versus the WFDB files read by the `rdsamp` function of WFDB. All 16-bit WFDB files had an SNR of above 60 dB per channel, as compared to the original MATLAB files. Although 60 dB is fully acceptable for most applications, researchers seeking double-precision floating point accuracy are advised to use the MATLAB files.

The description of the corresponding physical activities and the unique IDs of the participants are provided in the spreadsheet `ECGPCGSpreadsheet.csv`, which accompanies the dataset. For basic heart-rate extraction and analysis from the ECG and PCG channels a sample MATLAB script `TestHeartRateCalculation.m` is also provided online. Additional source codes for analyzing this data are available in the Open-Source Electrophysiological Toolbox (OSET) [4].

## IV. Analysis

The gathered database can be used for various ECG and PCG analysis under physical activities. For proof of concept, we present only some of the basic analysis, which can be performed on the heart-rate time-series obtained from the simultaneous ECG and PCG. The hypothesis is that while the heart-rate time-series obtained from the ECG and PCG are coarsely identical, there are “micro-deviations” between the two time-series, which are rich in information. In fact, apart from the methodologically different algorithms used for heart-rate calculation from the ECG and PCG time-series (due to the difference in their time and frequency domain specifications), the micro-deviations between the two time-series reflect the differences and the minor time-lags between the electrical and mechanical functions of the heart. These minor deviations are not random, and vary from case to case and under physical workloads. In order to assess this hypothesis, the signal processing steps shown in Fig. 4 were applied to the ECG and PCG channels. The outputs of the processing scheme in Fig. 4 are the heart-rate time-series. Each stage of the processing units is detailed in the sequel.

**Fig. 4.**
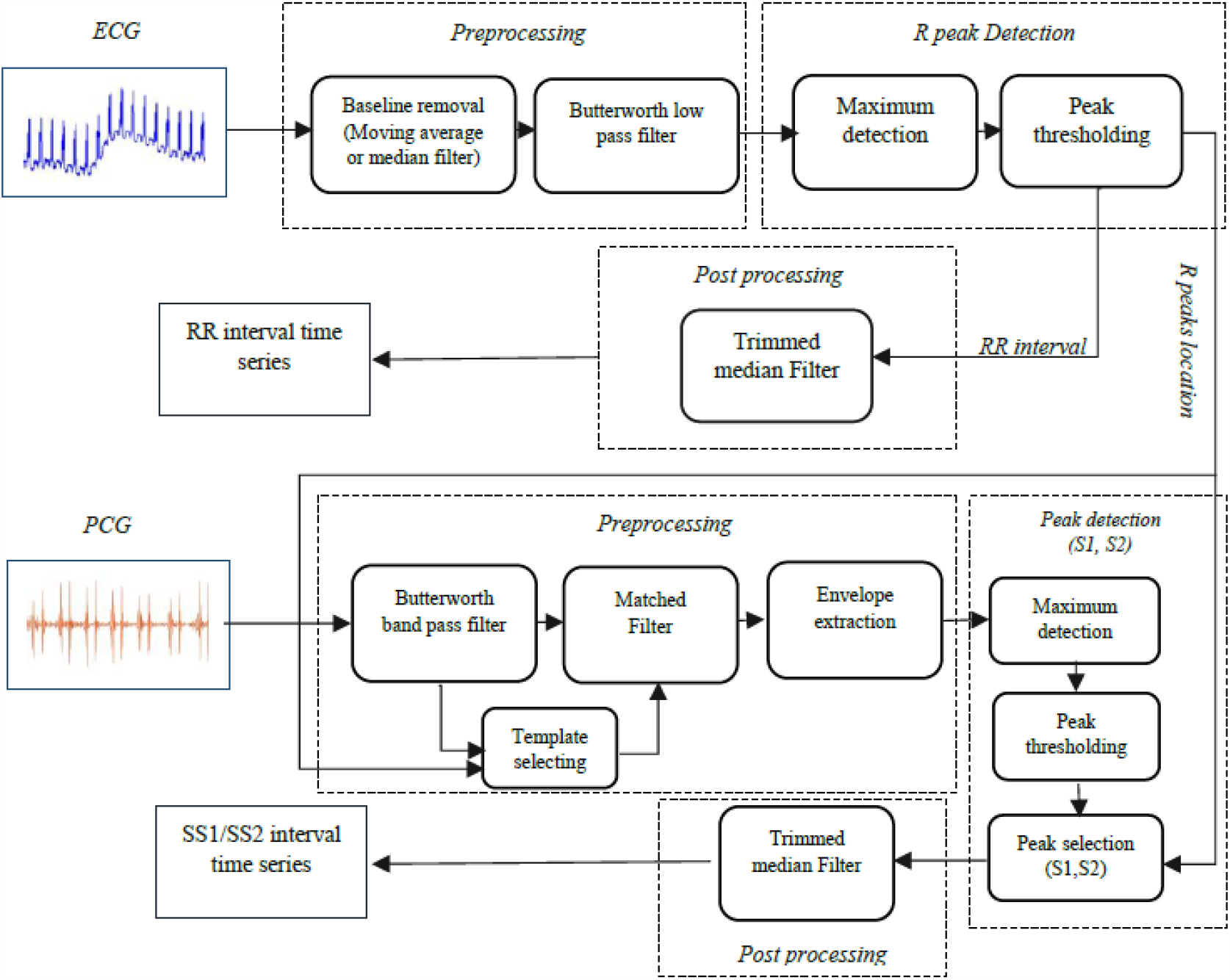
Signal processing block diagram for heart-rate calculation from ECG and PCG signals

### A. Pre-processing

For the ECG channels, a cascade of a moving median filter of length 0.6 s, and a moving average filter of length 0.3 s are used to remove the baseline wander. This combination has been previously shown to be very effective and robust for ECG baseline wander removal. As an acoustic signal, the PCG does not physically have a baseline drift and therefore does not require baseline removal. However, in order to keep the ECG and PCG synchronous, the delay of the two-stage baseline wander removal filters in the ECG signal path (which is half the total window lengths, or 0.45 s) is added to the PCG signal path, by zero-padding the beginning of the PCG signals.

Since the signals have been acquired indoor, the powerline noise (50 Hz in this dataset) was inevitable and appeared in variable amplitude throughout the thirty minute acquisition sessions. As specified in the dataset spreadsheet, the power-line has in cases appeared as bust noise of short duration. Therefore, a fixed notch filter was insufficient for powerline cancellation in these cases. Instead, an adaptive Kalman notch filter developed in [9] was used, which adapts the notch filter’s quality-factor (Q-factor) over time, depending on the level of powerline interference. The source codes for this notch filter are online available in the open-source electrophysiological toolbox (OSET) [4].

### B. ECG-based heart-rate calculation

A modified version of the Pan-Tompkins algorithm was used for R-peak detection from the ECG channels. Accordingly, a bandpass filter with a passband between 10 Hz and 40 Hz was applied to the ECG. Next, the signal energy envelope was calculated over a sliding window of length 75 ms. The R-peaks were obtained by local peak detection of the energy envelopes over windows of 0.5 s. This R-peak detection scheme was very effective for the current database and was approved by visual inspection of the R-peak locations on the ECG signals. Despite its accuracy, due to the physical activity of the subjects, a small fraction of the R-peaks were mis-detected or dropped. These cases were identified by studying the heart-rate time-series trend and by detecting outlier peaks in the RR-interval sequence. In these cases, the misdetected R-peaks were manually corrected by an expert analyzer during post-processing.

### C. PCG-based heart-rate calculation

ECG peak detection algorithms are directly applicable to the PCG. A major advantage of simultaneous ECG and PCG acquisition is that the R-peaks can be obtained from the ECG and used as a reference for beat segmentation and detection of the PCG components (S1, S2, etc.), which are otherwise not trivial to detect. For this study, the ECG R-peaks were first estimated from the ECG and used as initial reference points for estimating the S1 and S2 segments of the PCG. These major PCG components are characterized by bumpy shapes modulated over rather narrow band oscillatory waves. Therefore, contrary to the ECG, the local peaks of the PCG are not good indicators of the beat point. Instead, the energy envelopes in the S1 and S2 frequency bands are more robust indicators of the peak points of these events.

A typical PCG and its time-frequency representation is shown in Fig. 6. Accordingly, the major Frequency components, for both first heart sound (S1) and second heart sound (S2) are below 200 Hz. This property was approved through visual inspection of all data files. Therefore, for PCG-based beat detection, a Butterworth bandpass filter with 35 Hz-200 Hz passband frequency was applied to the PCG. The resulting signal was used to calculate the local energy envelope peaks, located after each ECG-based R-peak. The first and second local peaks in the PCG envelope, which occurred between successive R-peaks were assigned as the S1 and S2 components, respectively (cf. Fig. 5).

**Fig. 5.**
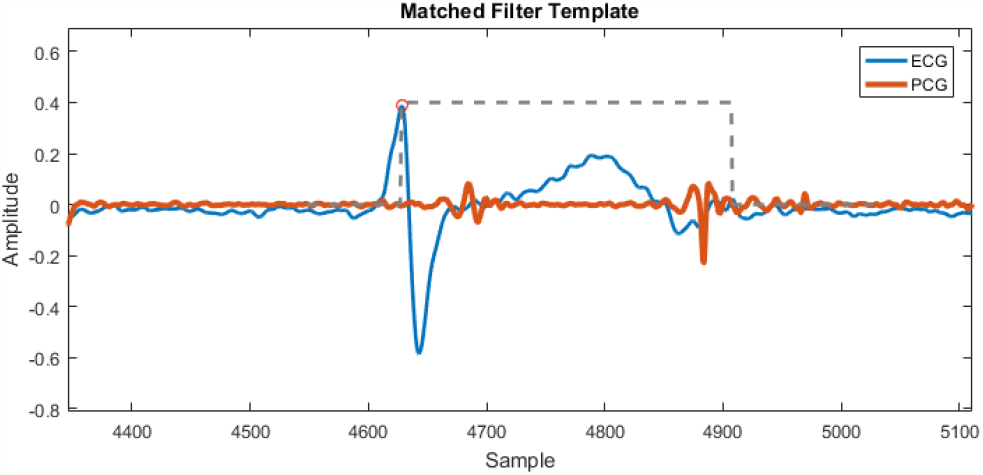
Matched filter template selection using R peak location

**Fig. 6.**
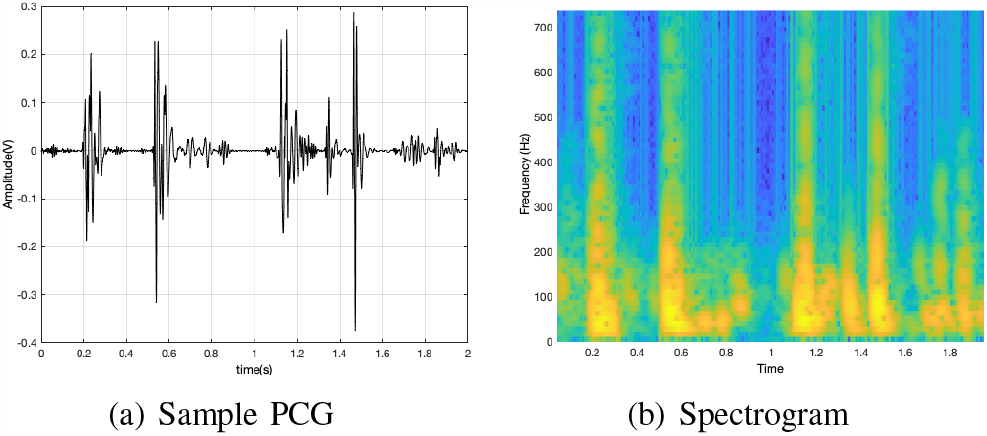
(a) Sample PCG and (b) the time-frequency domain of the PCG

### D. ECG versus PCG Heart-Rate Parameters

Having the R-peaks from the ECG and the S1 and S2 peaks of the PCG, the following time intervals were calculated per record: 1) R-R: from one R-peak to the next R-peak, 2) S1-S1: from one S1-peak to the next S1-peak, 3) S2-S2: from one S2-peak to the next S2-peak, 4) R-S1: from one R-peak to the S1-peak of the same beat, 5) R-S2: from one R-peak to the S2-peak of the same beat, 6) S1-S2: from one S1-peak to the S2-peak of the same beat (systolic time interval), 7) S2-S1: from one S2-peak to the S1-peak of the next beat (diastolic time interval).

An example of the resulting time sequences is shown in Fig. 7, for one of the stationary biking samples of our dataset.

**Fig. 7.**
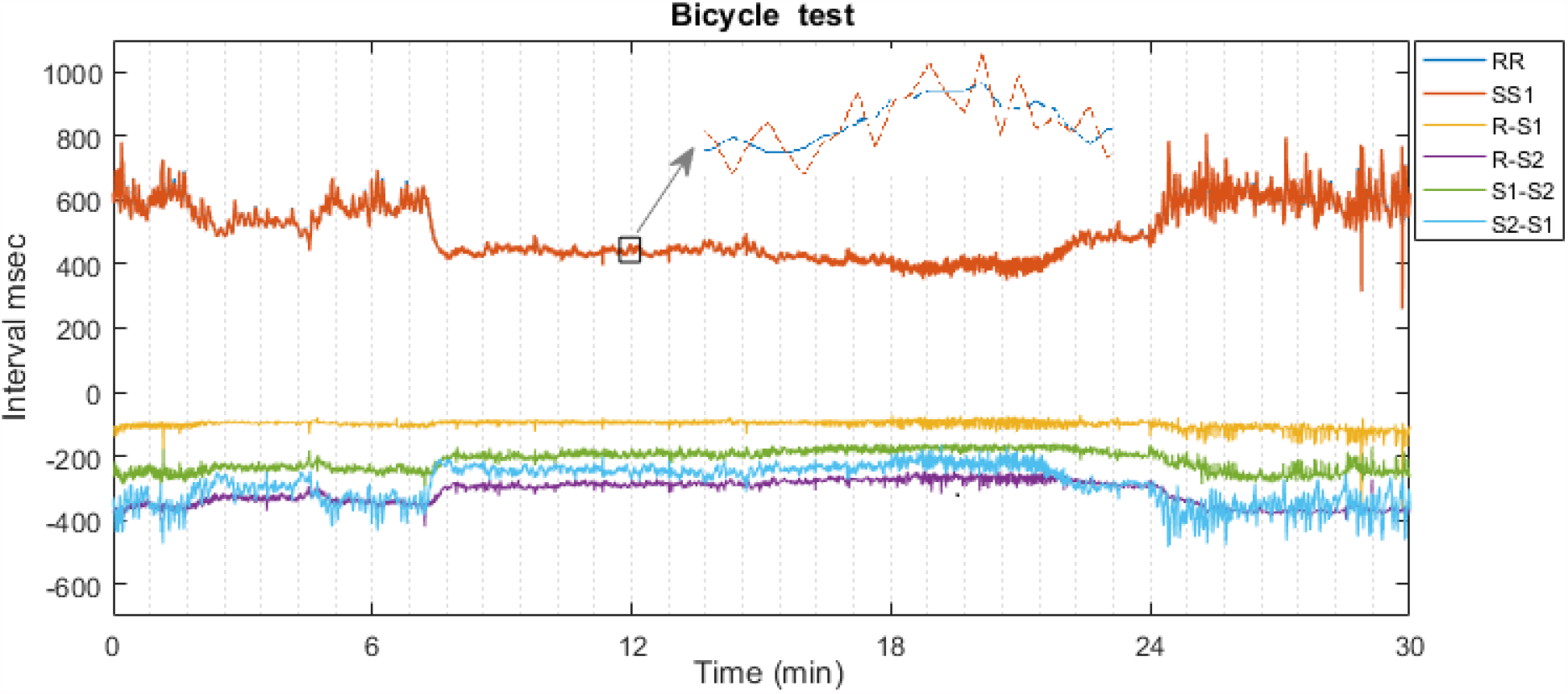
ECG and PCG based event time intervals of a subject pedaling on a stationary bicycle

The power spectrum of a typical heart rate variability (HRV) time-series calculated from the R-R intervals (from the ECG) and the S1-S1 intervals (from the PCG) are shown in Fig. 8.

As a sample nonlinear analysis the approximate entropy of the HRV time-series were calculated and compared for a subject at rest, as shown in Fig. 9. Accordingly, the time-series of the ECG-based HRV for smaller values of the tolerance parameter (*r*) has a higher entropy than the time series of the corresponding PCG.

**Fig. 8.**
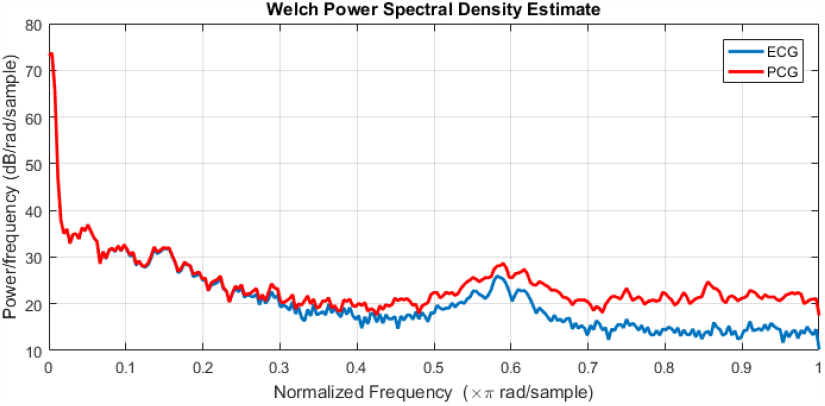
Welch power spectral density of HRV of a subject at rest mode.

**Fig. 9.**
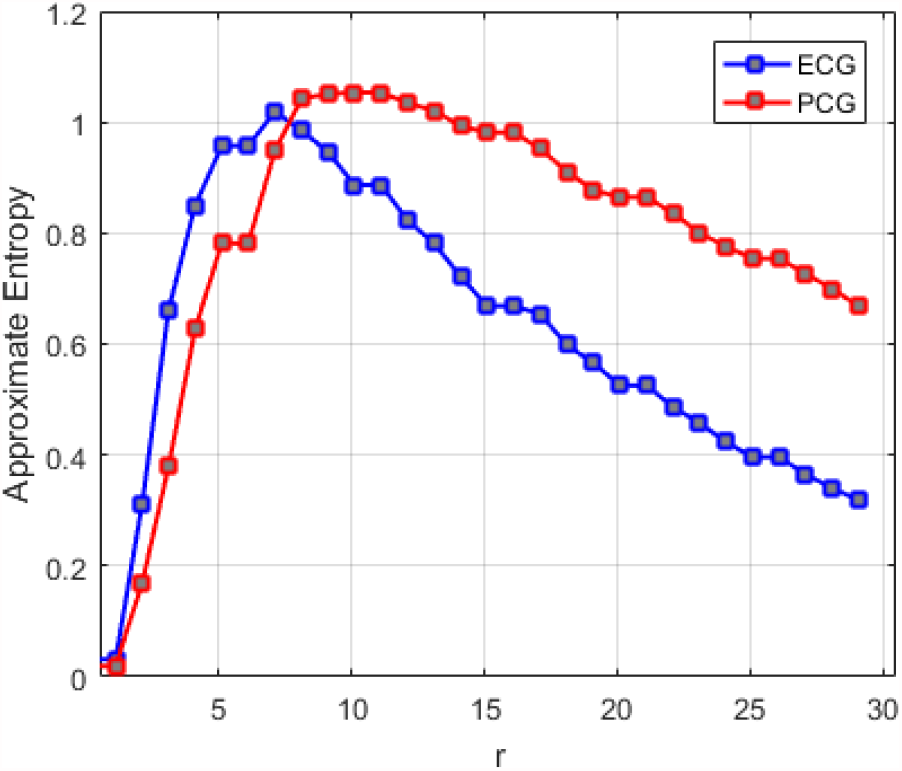
Approximate entropy of ECG/PCG HRV versus the *r* parameter

## V. Discussion

This current dataset is useful for simultaneous multimodal analysis of ECG and PCG, as it provides interesting insights into the inter-relationship between the mechanical and electrical mechanisms of the heart, under rest and physical activity. Only as proof of concept, the dataset was used to compare the heart-rate time-series and the heart-rate variabilities (HRVs) of the subjects during the aforementioned stress-test. It was shown that while the overall trend of the heart-rate time-series obtained from the ECG and PCG are exactly identical (as expected), there are considerable “micro-variations” between them, which reflects the differences between the electrical and mechanical functions of the heart during different levels of physical activity. Specifically, as the heart-rate increases and the RR-intervals of the ECG become shorter, the differences between the R-peaks and the first and second heart sounds (namely S1 and S2) obtained from the PCG channels do not scale at the same rate. There are also notable differences between stochastic features such as the sample entropy of the ECG- and PCG-driven heart-rate time-series.

Another application for the dataset is to use the simultaneous ECG and PCG channels to develop mathematical PCG models for generating synthetic signals. Previous research in electrocardiography, have shown how synthetic ECG generators [10], can be used to develop algorithms for Bayesian filtering and parameter extraction from highly noisy ECG recordings [11]. The current dataset can help researchers to develop similar algorithms for de-noising and automatic parameter extraction from the PCG.

As noted in the data description, the three channels PCG2, AUX1, and AUX2 (which are available for some of the records), are mostly very weak in amplitude (at quantization noise level). However, through visual inspection and by listening to these audio channels, it is noticed that they have captured some of the electronic device noises and the weak background sounds in the environment. Therefore, although they are not useful for direct utilization, researchers interested in the signal processing aspects of the dataset might find these channels useful for designing adaptive noise cancellers or multichannel blind and semi-blind source separation algorithms.

Due to the type of available chest braces used in this study and the anatomical points selected for better auscultation of the heart sounds, the setup was found to be inappropriate for females during stress tests (multiple attempts in recording signals from female volunteers were unsuccessful). This issue is addressed in future versions of our design, by developing a customized chest brace that would be comfortable for stress tests, while fixing the stethoscope and ECG leads in place.

## VI. Conclusion and Future Work

The methodological aspects of the current research can be extended from various aspects. We can for example study the mutual information shared between the ECG and the PCG, and the unique information through each modality, by using time and frequency domain features. Apparently, the mutual/unique information between the two modalities are extensively rich in information and may be used to study the micro-variations between the electrical and mechanical mechanisms of the heart. We can also develop a Kalman filter based heart-rate tracker, which uses the local peaks of the ECG and the PCG as complementary measurements, similar to the approach adopted in [12].

## Acknowledgement

The database was acquired as part of a master’s thesis in Biomedical Engineering at the School of Electrical and Computer Engineering of Shiraz University, Shiraz, Iran between 2016 and 2018.

## Footnotes

1 Note that while the ECG contains diagnostic information in a range of 0.7 Hz to 150 Hz, according to the IEC standard [5], if an ECG device is used for ST-segment measurements, a lower cutoff frequency of 0.05 Hz would be required.

